# Subicular spatial codes arise from predictive mapping

**DOI:** 10.1101/2024.11.06.622306

**Authors:** Lauren Bennett, William de Cothi, Laurenz Muessig, Fábio R Rodrigues, Francesca Cacucci, Tom J Wills, Yanjun Sun, Lisa M Giocomo, Colin Lever, Steven Poulter, Caswell Barry

## Abstract

The successor representation has emerged as a powerful model for understanding mammalian navigation and memory; explaining the spatial coding properties of hippocampal place cells and entorhinal grid cells. However, the diverse spatial responses of subicular neurons, the primary output of the hippocampus, have eluded a unified account. Here, we demonstrate that incorporating rodent behavioural biases into the successor representation successfully reproduces the heterogeneous activity patterns of subicular neurons. This framework accounts for the emergence of boundary and corner cells - neuronal types absent in upstream hippocampal regions. We provide evidence that subicular firing patterns are more accurately described by the successor representation than a purely spatial or boundary vector cell model of subiculum. Our work reveals a temporal hierarchy in hippocampal-subicular processing, with subiculum encoding predictive representations over longer time horizons than CA1, capturing extended behavioural patterns and environmental affordances.

## Introduction

The hippocampal formation is intimately linked to episodic memory and spatial cognition^1–3^ - functions believed to rely on distinct populations of spatially modulated neurons. Most notably, these include CA3/1 place cells^4^ and medial entorhinal cortex grid cells^5^, as well as several other cell-types distributed through the hippocampus and associated regions^6–9^. Collectively, these neurons, which constitute a representation of self-location, are held to form a cognitive map^10^ that supports flexible navigation and reasoning in both physical and abstract spaces^11,12^.

Compared to CA3/1 and entorhinal cortex, subiculum — the primary output structure of the hippocampus — has received comparatively little attention. This oversight is surprising given its computational potential: subiculum contains a larger number of principal neurons with more diverse morphology than CA3 and CA1 combined^13^, coupled with extensive recurrent connectivity^14,15^. Moreover, subiculum holds a privileged position in the connectome, with dense projections to an extensive cortical and subcortical network that includes anterior thalamic nuclei, retrosplenial, medial prefrontal, and entorhinal cortices^13,16,17^.

To a certain extent, this relative neglect can be attributed to the absence of a single clear computational role for subiculum. Subicular principal neurons exhibit a diverse menagerie of spatial responses^6^, with authors often choosing to emphasise one distinct aspect of their observed activity. Amongst these, boundary vector cells are perhaps the best known. Initially theorised as an allocentric input to place cells^18^, neurons matching this description were subsequently identified in subiculum^19,20^. However, these cells lie downstream of primary CA3/1 projections^17^, raising the possibility that they may be derived from place cell activity, rather than contributing to it. Boundary vector cells are characterised by elongated firing fields that run parallel to boundaries and generally align with the surrounding environmental geometry^20,21^. A subsection of these fields also generate ‘trace’ responses that persist after barrier removal^8^. These characteristics have been construed as demonstrating a potential role for subiculum in representing and recalling environmental geometry and physical affordances^8,19,20,22^.

Further studies have highlighted the role of subiculum in representing movements and trajectory sequences. For example, subicular neurons have been reported to represent the current axis of travel^23^, as well as composite signals of place and head orientation^24^, specific trajectories^25^, running speed^25,26^, and head direction^23,26^. Adding to this complexity, Sun *et al*.^9^ recently identified subicular neurons that represent corners — both concave and convex — within an environment.

Unlike subiculum, hippocampal place cells have attracted considerable attention from theoreticians proposing normative and mechanistic models^27–30^. While hippocampal representations extend beyond simple spatial coding—encompassing theta sequences^31^, replay^32,33^, remapping^34,35^, multi-scale representations^36,37^, and non-spatial variables^12^—the core place cell phenomenon has proven amenable to unified computational accounts. Recently, predictive algorithms have emerged as a powerful framework for understanding hippocampal function and the generation of neural responses in associated regions^38–43^. In particular, the successor representation, which calculates expected future state occupancies^44^, accounts for how place fields are shaped by both environmental boundaries and animal behaviour^39,45^. Importantly, the successor model consolidates evidence that agent behaviour greatly influences spatial representations^46–48^, and explains navigational and search biases observed in humans and rodents performing spatial tasks^49,50^. In contrast, the diverse spatial phenotypes observed in subiculum—boundary cells, corner cells, trace responses, and trajectory-dependent coding—have resisted such unification, appearing instead as a heterogeneous collection of distinct computational roles.

While the successor representation can model spatial responses as long-run predictions over discrete locations^39^, it can also be learnt over amalgams of sensory information termed ‘features’ that correspond to a more plausible neural representation of state^45,51^. Building on this, recent work has shown that biological computations within the hippocampus are sufficient to approximate successor learning^52–54^. Thus, spike-timing-dependent plasticity^55^ (STDP) over sequences of place cell firing, ordered by theta sequences, can sweep into subiculum^28,56,57^ and provide a physiological mechanism that rapidly encodes transitions between fields.

Although in previous work the successor representation has been deployed as a model of CA3/1, we here propose a reconceptualisation based on the position of the hippocampus atop a multi-modal sensory hierarchy. Thus, place cell responses can be viewed as generalised amalgams of information from diverse modalities^37,58,59^ – corresponding to neural representations of spatial or non-spatial states depending on the available sensory input. We posit that STDP acting on theta sweeps over these states^52–54^, or equivalent computations^52–54^, yields successor features in the downstream subiculum that align with its unique anatomical and physiological properties. Furthermore, because rodents exhibit stereotyped behaviours, such as thigmotaxic running and corner-dwelling, we hypothesise that successor features based on biologically realistic trajectories, using place cell activations as basis features, will more closely match the statistics, features, and appearance of subicular representations than those found in other hippocampal regions.

Here, we test this proposal by comparing predictions from the successor representation model to two electrophysiological and one calcium imaging dataset recorded from rodent subiculum^8,9,21^ and hippocampus^60,61^. We find that successor features, derived from place cell basis features and trained on real rodent trajectories, closely resemble subicular spatial responses. Specifically, this simple model generates corner responses^9^ and border activations that correspond to boundary vector cells^19,20^. We demonstrate that the boundary responses emerging from the successor representation framework better match the population statistics of subicular boundary cells than predictions derived from the boundary vector cell model^18^. Furthermore, we use representational similarity analysis^62,63^ (RSA) to quantify that a successor representation model based on rodent trajectories fits a composite dataset comprised of three subicular experiments better than alternative models. Crucially, we identify distinct temporal horizons for hippocampal versus subicular predictive representations, with subiculum integrating information over substantially longer timescales than CA1. Collectively, these findings support a computational account of subicular function - positioning it as a predictive map of the environment, operating at extended temporal horizons, derived from hippocampal inputs.

## Results

To investigate whether subiculum implements a successor representation over hippocampal states, we modelled CA1 as a set of spatially-modulated ‘basis features’, *ϕ*(*s*_*t*_), that resemble the observed statistics of hippocampal place cells^37^. Thus, in a rectangular environment, these sparse basis features vary with location, *s*_*t*_, and are instantiated as thresholded 2D Gaussian fields (see Methods) that increase in size towards the centre of the environment and compress orthogonal to boundaries (Fig.1a; key results replicated with symmetric, uniform sized Gaussian fields in Supplementary Information Figure S1). The longest distance rodent trajectory from each of the three datasets^8,9,21^ was down-sampled to 10-12 Hz (Fig.1b; Methods) and the successor matrix, *M*, (Fig.1c) was updated at each time point according to the temporal-difference learning rule (derived in Appendix):

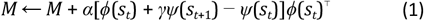

where:

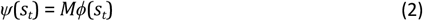

are the corresponding successor features.

**Fig.1.**
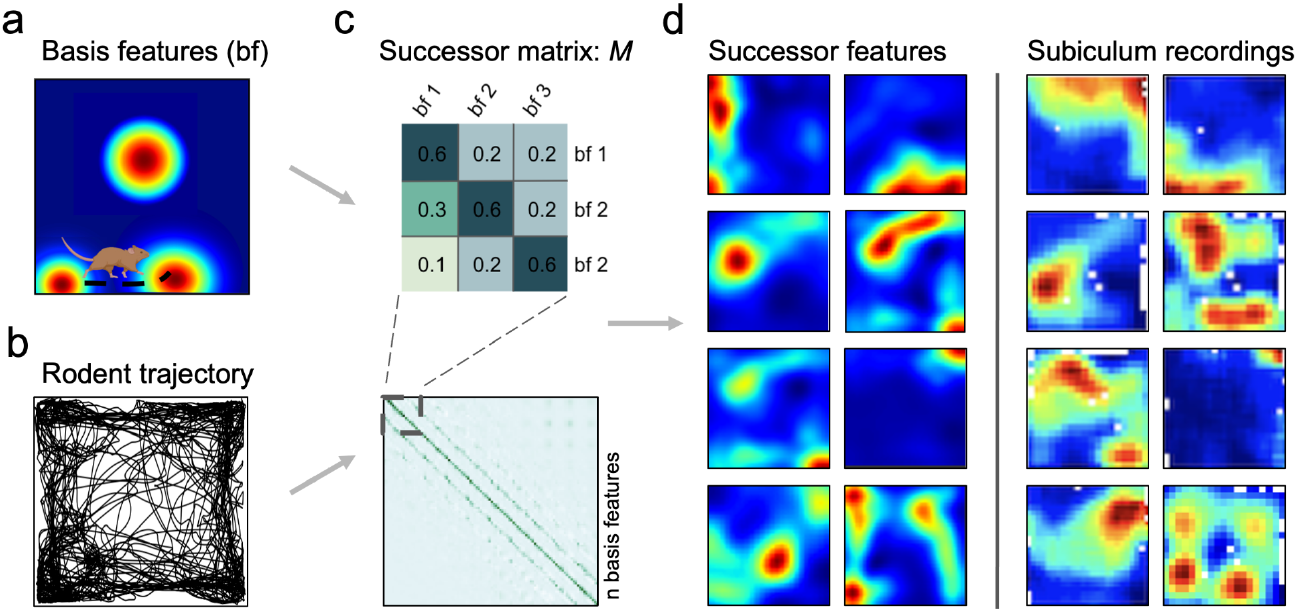
Model pipeline. (**a**) Biological basis features (n=400 for each of the three environmental sizes) were sampled along (**b**) real rodent trajectories in square environments in order to learn the successor matrix, *M* (**c**). A representative single trial trajectory in a 100cm square (**b**) highlights rodent thigmotaxis and preference for corner dwelling; with 85.7% occupancy in the perimeter versus the equally-sized inner area (mean distance of animal to nearest wall=11.8cm). Once trained, successor features are formed by the matrix multiplication of *M* with the population vector of basis features. (**d**) Exemplar successor features generated from rodent trajectories (left columns) closely resemble rodent subicular responses (right columns), here recorded by Sun *et al*. 2024.

The population activity of successor features at a given time, *ψ*(*s*_*t*_), constitutes a predictive code that collectively captures long-run expectations about upcoming spatial states. Thus, these expectations recapitulate rodent behavioural biases, such as a proclivity to run along boundaries (thigmotaxis) and dwell near corners. The resultant successor features extend along walls and into corners (Fig.1d left); ultimately resembling the firing patterns of subicular neurons (Fig.1d right).

### Boundary responses

We first set out to investigate whether successor features provide a better account of subicular boundary responses than prior theories - specifically the boundary vector cell model^18–20^. Boundary-responsive neurons in subiculum (Fig.2a) are explained in the boundary vector cell model as having receptive fields that are tuned at certain distances and directions to environmental boundaries^18^ (Fig.2b). Conversely, the successor feature model (Fig.2c) proposes that boundary responses arise from predictable patterns in animal movement - particularly thigmotaxic running alongside walls^64^. To quantify these behavioural biases, we segmented each environment into a 3×3 grid (Fig.2d top) and calculated the polar histogram of heading directions in each segment (Fig.2d bottom). Trajectory heading directions in segments adjacent to boundaries were significantly less heterogeneous than those in the central portion of the environment (Fig.2e; pairwise t-test between Kullback-Leibler divergence from uniform distribution for the longest distance trajectory from each trajectory used to generate successor features, t(2)=-9.97, p=0.010). While the boundary vector cell model describes individual cell tuning^18,19,45^, it does not explain why boundary cell populations show environment-specific organisation^21^. We hypothesised that the successor feature model naturally reproduces these geometry-dependent distributions due to behaviour-environment interactions. Indeed, in light of these pronounced behavioural biases, the boundary vector cell and successor feature models make competing and testable predictions about the morphology of boundary responses in subiculum. Specifically, canonical implementations of the boundary vector cell model typically sample parameters uniformly, generating populations with vector responses that evenly span all directional tunings and encompass distance tunings sufficient to cover the local environment^18,19,45^, in order to provide a comprehensive and boundary-centred representation of allocentric space. Conversely, rodents’ proclivity to run adjacent and parallel to environmental boundaries means that the successor feature model predicts subicular responses that align proximal to environmental walls.

**Fig.2.**
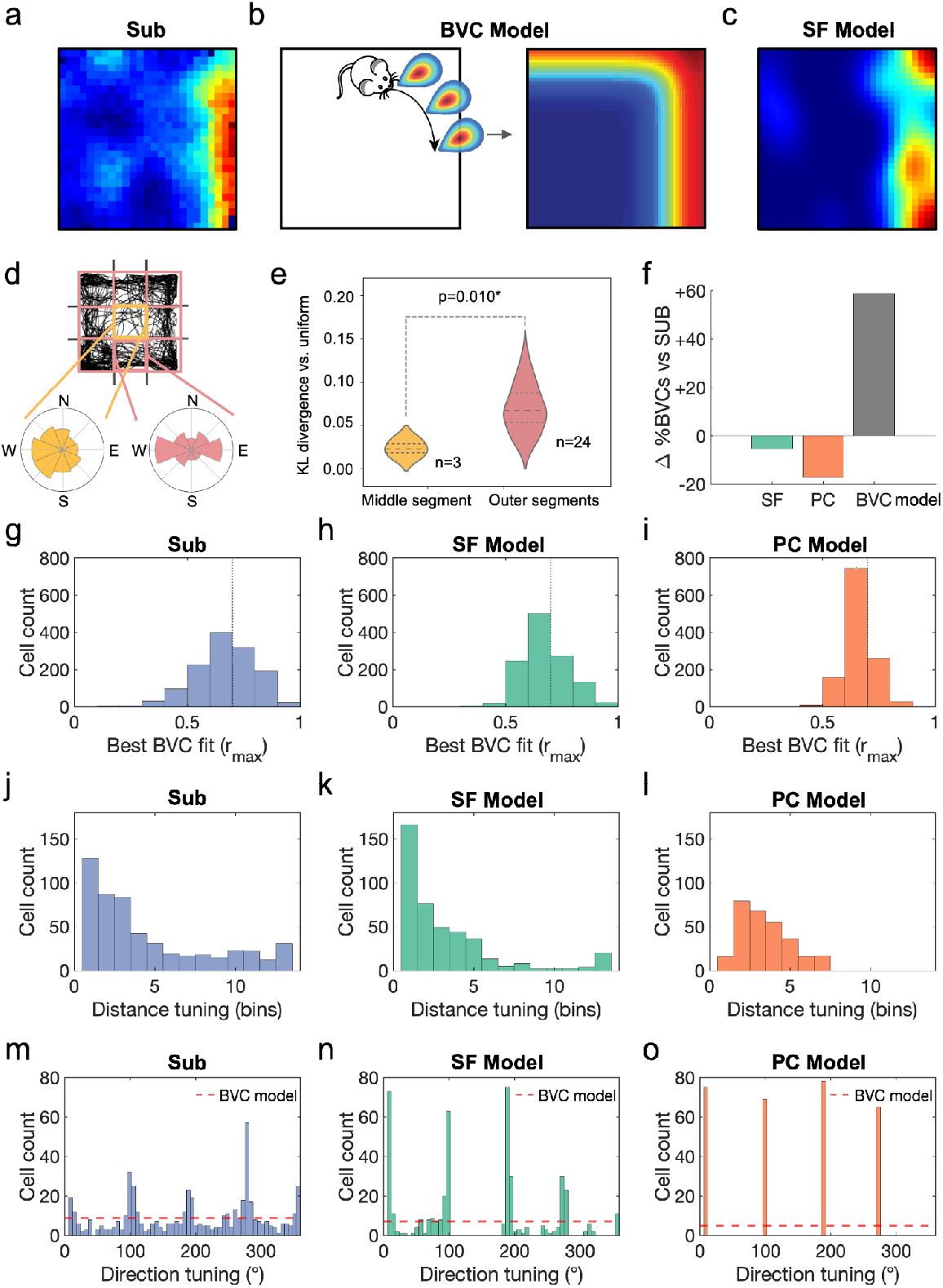
Successor features produce subicular boundary representations. (**a**) Boundary-responsive cells in rodent subiculum have previously been explained by (**b**) the boundary vector cell model as having receptive fields tuned to environmental boundaries at certain directions and distances. Conversely, the successor feature model (**c**) predicts that behavioural biases near walls produce similar boundary responses in the resulting successor features. This is due to (**d**) trajectory heading directions being more heterogeneous in the central portion of the environment than at the perimeter adjacent to walls, where trajectories are constrained both directionally and by anxiety behaviours such as thigmotaxis, measured by (**e**) the Kullback-Leibler divergence of trajectory headings vs uniform distribution (pairwise t-test middle vs outer segments, using the longest distance trajectory from each dataset, t(2)=-9.97, p=0.010). (**f**) Comparing the proportions of boundary cells observed the subicular data (Sub) to models of subicular function – successor features (SF), a Gaussian place cell control model (PC), and the boundary vector cell model (BVC) – identifies the successor feature model as closest to capturing the subicular data (−5.5% discrepancy), with the place cell control and BVC models respectively under- and overestimating the proportion of subicular boundary cells (−17.2% and +58.9%). Fitting the boundary vector cell (BVC) model to subicular data (SUB), successor features (SF) and a Gaussian place cell control model (PC) revealed model fits that were most similar between (**g**) the subiculum data and (**h**) the successor feature model (t-test following Fisher z-transform: t(2483)=0.057,p=0.955; dotted line at r_max_=0.7 indicates threshold criteria for a boundary cell), producing significantly better fits than (**i**) the place cell control model (SUB vs PC, t(2483)=5.67, p<0.001; SF vs PC, t(2483)=6.54, p<0.001). (**j**) Subicular boundary cells were found to over-represent boundary responses proximal to environmental walls, better matching (**k**) the successor feature model than (**l**) the place cell control (pairwise t-test on absolute probability density residuals between SUB-SF vs SUB-PC, t(12)=3.672, p=0.003). Crucially, (**m**) vectorial responses in subiculum were clustered around directions orthogonal to the walls in a square arena (quadrupled, wrapped directional tunings to test four-fold symmetrical clustering: Rayleigh test for non-uniformity, v=90.9, p<0.001), as predicted by (**n**) the successor feature model (quadrupled, wrapped direction tunings, Rayleigh test for non-uniformity, v=229, p<0.001; Watson-Williams test of circular means, SUB vs SF, F_1,953_=0.54, p=0.462), while the canonical boundary vector cell model predicts evenly-distributed tunings^18,19^ (dashed line) and (**o**) the place cell control model only detects vectorial tunings perpendicular and slightly distal to boundaries. *p<0.05; **p<0.01;***p<0.001.

To test these two predictions, we adopted the procedure of Muessig *et al*.^21^ and fit the boundary vector cell model to subicular^8,9,21^ and successor feature data. We simulated n=3120 boundary vector cells in a square environment for each of the three rodent datasets^8,9,21^, using vectorial direction tunings spanning 0-354° (6° increments) and preferred distance tunings covering 4-52% of the environment’s dimension (4% increments). Following Muessig *et al*.^21^, we classified cells as boundary cells when the Pearson correlation between their rate map and the best-fitting boundary vector cell model rate map exceeded 0.7, using an exhaustive search procedure to maximise this correlation. We then compared the proportion of boundary cells in the empirical subicular recordings (n=1285 cells) to three models (Fig.2f): the successor feature model (n=1200, trained on the longest rodent trajectory from each dataset), a Gaussian place cell control (n=1200), and the boundary vector cell model itself (n=3120). The successor feature model most accurately captured the proportion of boundary cells observed in subiculum, with only a -5.5% discrepancy. In contrast, the place cell control underestimated boundary cells by 17.2%, while the boundary vector cell model overestimated by 58.9%. The best fitting correlation values to the boundary vector cell model were similar between the subicular data (Fig.2g) and the successor feature model (Fig.2h; t-test following Fisher z-transform: t(2483)=0.057, p=0.955) with both producing significantly better fits than the place cell control model (Fig.2i; SUB vs PC, t(2483)=5.67, p<0.001; SF vs PC, t(2483)=6.54, p<0.001).

In line with our previous work^21^, subicular boundary responses over-represented short-distance tunings (Fig.2j), and provided a better match to the successor feature model (Fig.2k) than the place cell control (Fig.2l; pairwise t-test on magnitude of probability density residuals between SUB-SF vs SUB-PC, t(12)=3.67, p=0.003). Further, following ^21^, subicular vectorial responses were clustered near directions orthogonal to environmental walls, such as the cardinal axes of a square arena (Fig.2m). We quantified this four-fold symmetrical clustering using the Rayleigh test on the quadrupled, wrapped directional tunings from the boundary vector cell model fits (Rayleigh test for non-uniformity, v=90.9, p<0.001). These characteristics were matched by the successor feature model (Fig.2n; Rayleigh test for non-uniformity, v=405, p<0.001; Watson-Williams test of circular means, SUB vs SF directional tunings, F_1,953_=0.541, p=0.462), where stereotyped behaviour along walls, coupled with more heterogeneous behaviour away from boundaries (Fig.2d,e), generated an absence of long-range and off-axis boundary responses. Conversely, the place cell control model was only matched to perpendicular vectorial tunings (Fig.2o; Rayleigh test for non-uniformity, v=287.0, p<0.001) and less proximal (Fig.2l) boundary distances.

Our analyses reveal two key properties of successor feature boundary responses that align with empirical observations. First, these responses show resistance to place cell remapping: while remapping in the place cell basis can propagate to some successor features (Fig.3a), the boundary code largely maintains stability despite moderate upstream changes (Fig.3b). This stability could explain the rapid formation and persistence of boundary responses in novel environments^8,20,24^. Second, when barriers are inserted into familiar environments, successor features naturally develop duplicated boundary fields on either side of the barrier (Fig.3c)—a phenomenon considered diagnostic of boundary vector cells and observed in both hippocampal^19,65^ and subicular recordings^8,19,20^. In our model, this duplication emerges when a small subset of place cell fields duplicate against the inserted wall^19,65^, with successor features showing higher duplication rates than their basis features (Fig.3d).

**Fig.3.**
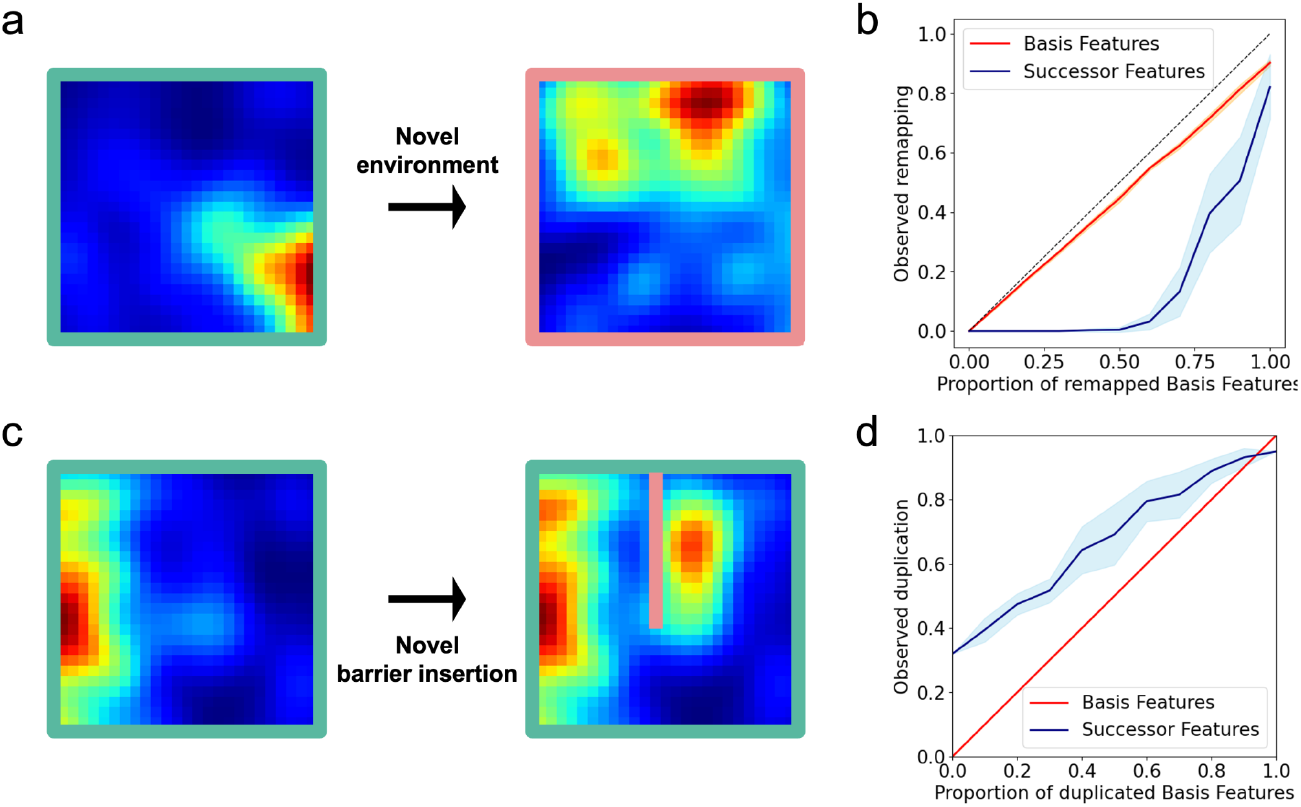
Successor feature code exhibits hallmarks of subicular representations: resistance to remapping and boundary field duplication. When animals enter novel environments, hippocampal place cells often remap. In our place cell basis feature model, this remapping (**a**) can propagate to some downstream successor features, but **(b**) the successor feature code largely resists these perturbations, maintaining stability despite moderate remapping in the underlying basis features (shaded regions indicate standard deviation calculated across 10 random seeds). This mirrors empirical observations where subicular spatial representations, including boundary vector cells, remap less frequently than hippocampal ones^8,20,24^. Similarly, when barriers are inserted into familiar environments, subicular boundary cells^8,19,20^ (and to a lesser extent hippocampal place cells^19,65^) duplicate their firing fields on either side of the barrier. Our model captures this phenomenon: (**c**) successor features derived from place cell basis features show field duplication, with (**d**) higher duplication rates in successor features compared to basis features. These patterns align well with reported differences between subicular and hippocampal spatial representations.

Notably, the successor feature model predicts that individual biases in animal trajectories will propagate into inhomogeneities in spatial fields. Indeed, such models have previously been used to explain the backwards skewing of CA1 place fields against the direction of travel on linear tracks^39,54,66^. In the successor feature model, this anticipatory skew occurs because earlier locations are predictive of an animal’s future position along the track. If subicular representations do arise from a successor framework learnt over CA1 basis features, they should also exhibit path-dependent skewing - a core property of successor features^39,54^. Specifically, we would expect subicular fields to exhibit greater path-dependent skewing between consecutive trials than CA1 place cells recorded in the same environment. We reasoned that this effect should be particularly evident in border-responsive neurons, given that rodents have stronger behavioural biases near environmental boundaries.

To test this proposition, we identified subicular cells with firing fields that were adjacent to boundaries in two consecutive trials (Fig.4a; peak firing <=5 to 6 bins from a wall, corresponding to <12.5cm for data from ^21^ (n=34 cells) and <10cm from ^9^ (n=113 cells)– data from ^8^ did not include consecutive trials without other manipulations). We averaged each cell’s main firing field onto its long axis (x or y) and calculated the change in its 1-dimensional centre of mass between trials as its skew (Fig.4b). Next, we quantified the bias of rodent runs through these fields, parallel to the cardinal walls (Fig.4c,d; see Methods). For those cells that were recorded over two consecutive trials (n=147 cells, 294 ratemaps), we found that subicular fields skewed significantly against the dominant axis of travel (Fig.4e; Spearman’s r(145)=-0.22, p=0.004). The skew of CA1 fields (n=105 cells, 210 ratemaps) recorded in the same environment was not significant (Fig.4f; Spearman’s r(103)=-0.11, p=0.13), likely because without specific goal-location manipulations^67,68^ behaviour in a 2-dimensional environment is less biassed and directionally constrained than on an linearised track^66^.

**Fig.4.**
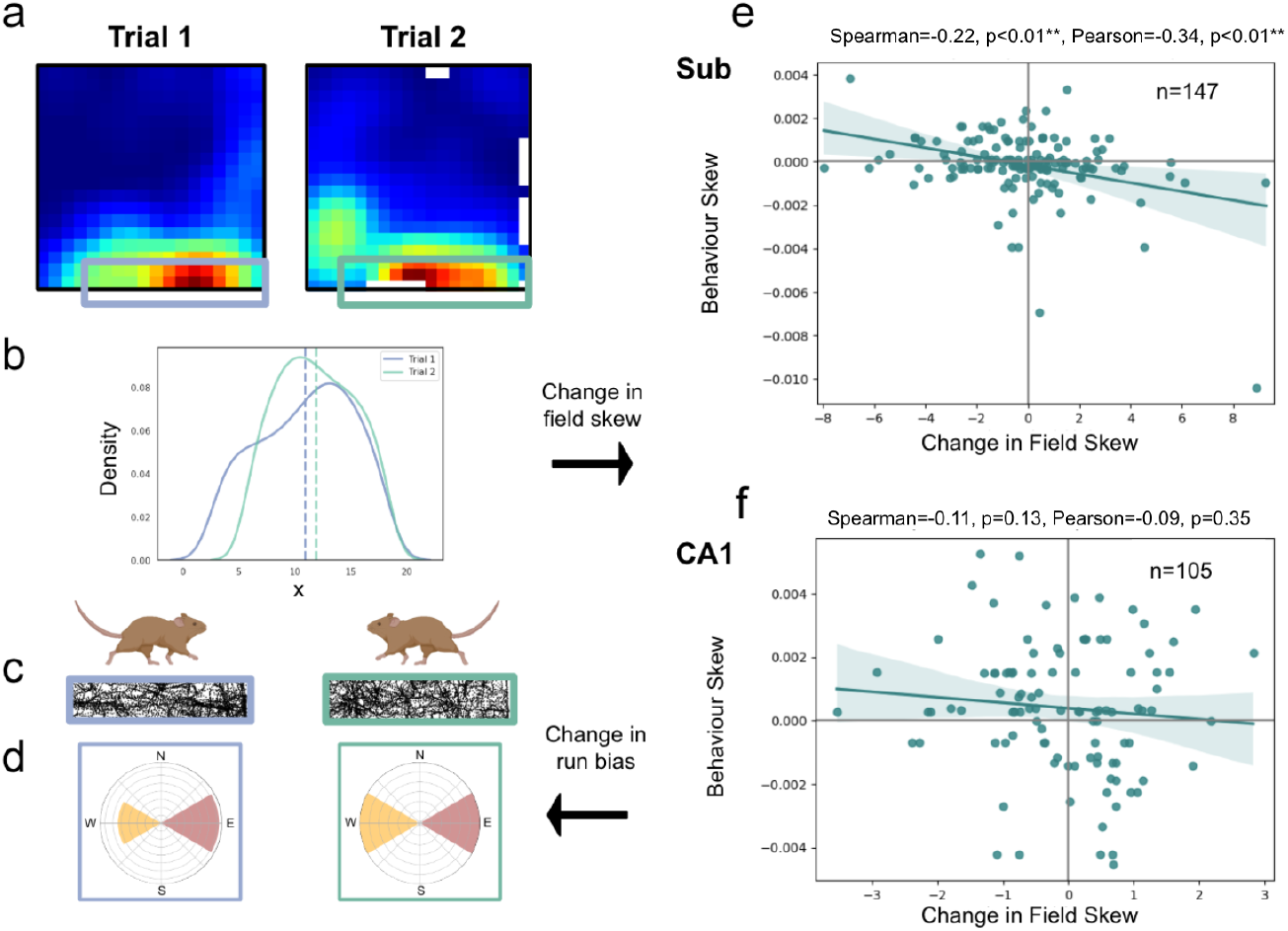
Boundary-tuned subicular responses are skewed by behavioural biases. The successor representation framework predicts that spatial firing fields will skew against an agent’s dominant axis of travel. To quantify this, (**a**) boundary-responsive CA1 and subicular fields were isolated and (**b**) averaged onto a single dimension, parallel to the adjacent wall. (**c**,**d**) The change in the directional bias of rodent runs through these fields was then regressed against the change in their centre of mass between consecutive trials. (**e**) Consistent with the predictive successor framework, subicular fields skewed significantly against rodents’ dominant axis of travel. (**f**) Notably, this backwards skew was not evident in corresponding CA1 fields. *p<0.05; **p<0.01;***p<0.001.

### Corner cells

Neurons that fire when an animal is present at an internal corner of the environment^9^ have recently been characterised as a unique subicular subclass, distinct from boundary-responsive cells (Fig.5). We hypothesised that these corner representations might also be accounted for by a successor representation model that recapitulates animals’ behavioural biases. Specifically, when rodents dwell in adjacent corners, linked by rapid runs along the intervening walls (Fig.5a-f), the resultant successor features integrate positional information over multiple corner locations.

**Fig.5.**
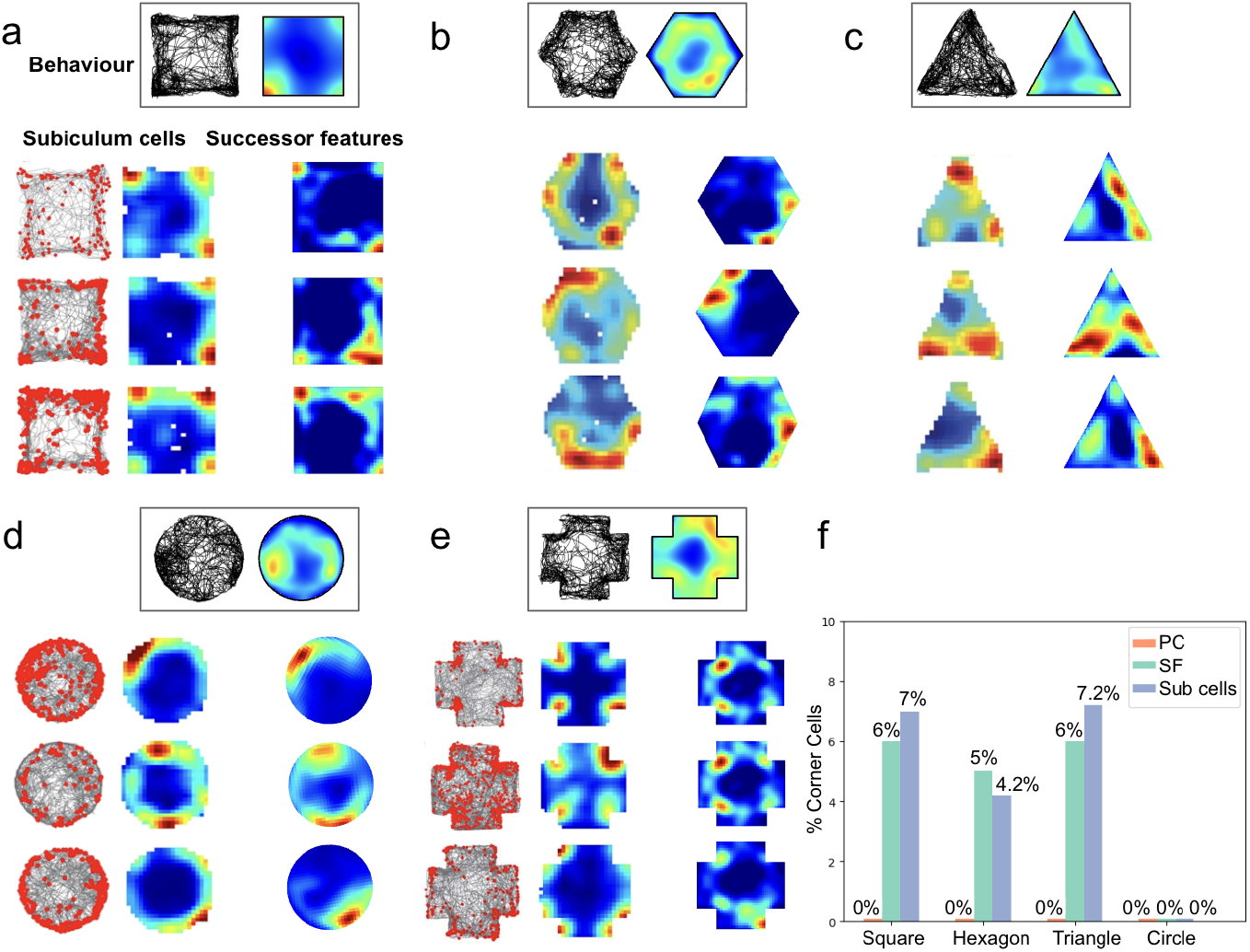
Successor features trained on real rodent trajectories exhibit subicular-like corner responses. Sun and colleagues recently characterised corner representations in subiculum as a distinct neural sub-class. (**a-e**) Top: rodent trajectories and positional heatmaps in square, hexagon, triangle, circle and cross environments. Behavioural paths highlight rodents’ preferences for wall-following and corner-dwelling. Bottom-left: subicular recordings demonstrating corner activity (data from ^9^). Bottom-right: successor features generated using real trajectories. (**f** Percentages of corner cells, defined using the ‘corner score’ metric^9^, were comparable between successor features and subicular recordings, across multiple environmental geometries. In contrast, negligible place field basis features were classified as corner cells. The proportion of corner cells in the cross environment was not reported in Sun *et al*.^9^.

In order to quantify corner representations in the successor representation model, we adopted Sun and colleague’s ‘corner score’ metric^9^. Across a range of different environmental geometries and using symmetric Gaussian basis features, the successor representation model generated a strikingly similar proportion of corner cells to those recorded empirically in subiculum. As before, we generated successor features for each geometry using the longest rodent trajectory and the same parameters (including *γ, α* and sampling rate) as in our other analyses.

For example, in a 100cm square environment, 6% of successor features were classified as corner cells, closely matching the 7% of subicular cells classified experimentally. Statistical comparison across geometries confirmed no significant difference between the successor model and experimental proportions (χ^2^(1, N=100)=0.20, p=0.905, Fig.5f; see Methods). In contrast, 0% of place field bases were classified as corner responsive, highlighting the specificity of the successor representation for generating corner cells.

### Population comparison

Next, to compare population-level representations, we employed representational similarity analysis (RSA) following Lee *et al*.^62,63,69,70^. For each ratemap (from both experimental recordings and model populations), we partitioned the spatial representation into a 3×3 grid and flattened each of the 9 segments into a vector. We then computed pairwise Pearson correlations between all 9 segment vectors within each ratemap, yielding a 9×9 similarity matrix per cell. These individual matrices were averaged within each population to generate a single 9×9 RSA matrix—one for each experimental dataset (subicular or CA1 recordings) and one for each corresponding model population (400 successor features trained on the longest trajectory from that dataset). Finally, we quantified model fit by calculating Pearson correlations between each experimental RSA matrix and its corresponding model RSA matrix (Fig. 6a). This approach captures the spatial structure of neural representations, as previously validated for place cell populations^63^.

**Fig.6.**
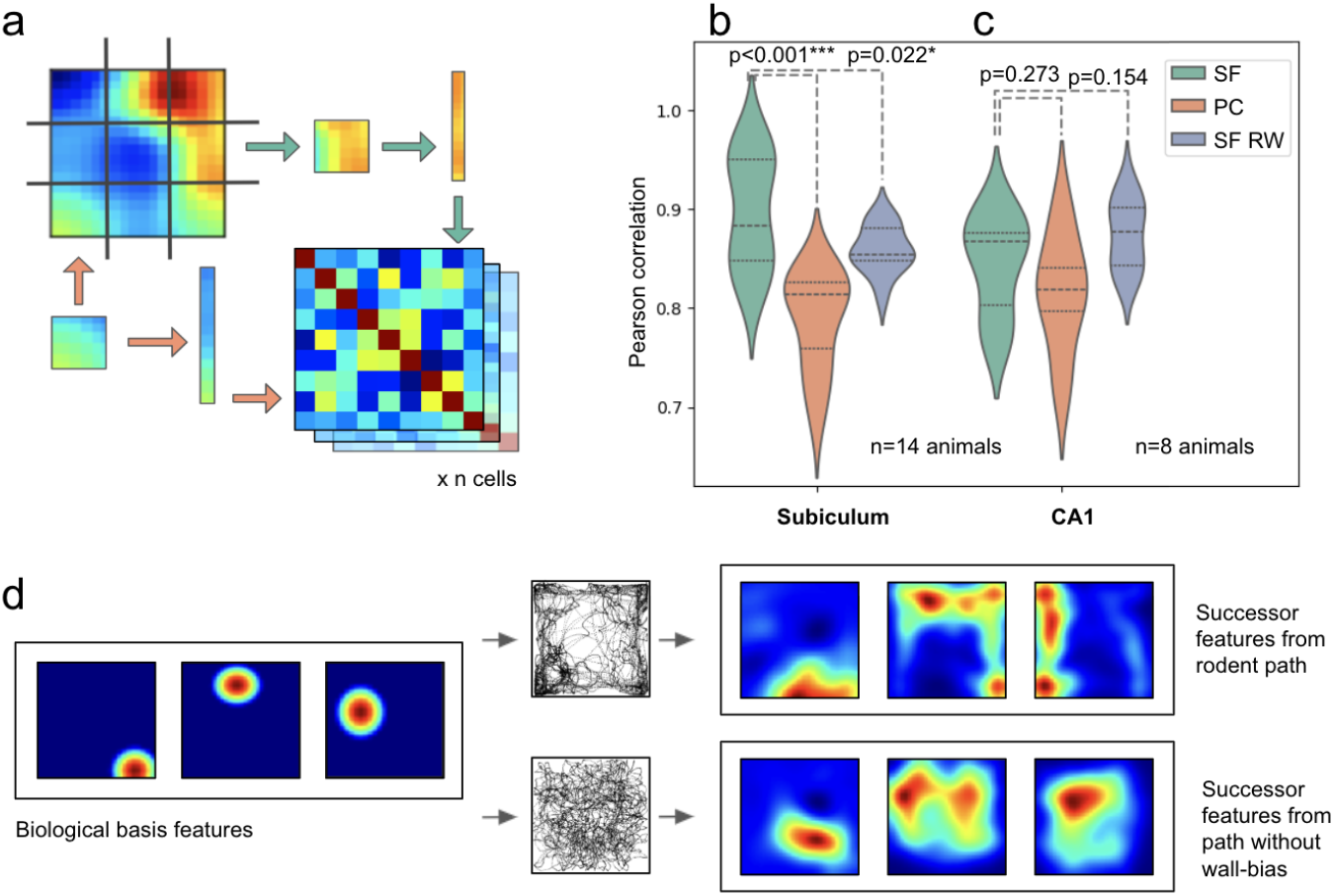
Subicular spatial responses are best described by a successor representation model trained using rodent trajectories. (**a**) Ratemaps from subicular recordings, successor features and place cell populations were each partitioned into nine sections, flattened, and cross-correlated. 9×9 correlation matrices were then averaged at either the rodent or rodent-trial level. (**b**) RSA correlations between populations of subicular cells to i) successor features based on rodent trajectories (SFs), ii) biological basis features modelled on place cells (PCs), and iii) successor features based on a biological random walk trajectory (SF RW). (**c**) Equivalent RSA results for a population of CA1 cells also recorded in the ^21^ environment are not better fit by SFs than SF RWs. Subicular responses, but not CA1 place fields, are fit better by successor features trained on real rodent trajectories. (**d**) Cartoon illustration of how wall-biassed behaviour, compared to a random walk, skews biologically inspired place cell basis features into elongated successor features, akin to subicular activations. *p<0.05; **p<0.01;***p<0.001.

Our analysis used subiculum data^8,9,21^ from 9 rats and 5 mice (9 to 366 cells per animal, n ratemaps=2425) and CA1 data recorded from 8 rats^60,61^ (10 to 78 cells per animal, n ratemaps=311) in square environments (box sides 30-100cm). We compared these condensed measures of biological population activity to similar matrices constructed for: i. Successor features built using the longest distance trajectory from any animal for each of the three datasets^8,9,21^ (SF); ii. Successor features trained on synthetic random walk paths^71^ with inertia but no tendency for wall-bias or thigmotaxis (SF RW) and; iii. Biological place cell basis features used to derive i and ii (PC).

Ultimately, we found that the successor representation model provided a better fit to subicular cells than competing models (Fig.6b; SF vs PC: 0.90 vs 0.79, t(13)=6.21, CI=(0.31,0.63), p<0.001; SF vs SF RW: 0.90 vs 0.86, t(13)=2.62, CI=(0.04,0.46), p=0.022) - results that highlight the importance of animals’ behavioural biases in shaping successor features. Conversely, there was no difference between the fit of the three models when applied to CA1 recordings (Fig.6c; SF vs PC: 0.84 vs 0.82, t(7)=1.17, CI=(−0.09,0.27), p=0.273; SF vs SF RW: 0.84 vs 0.87, t(7)=-1.55, CI=(−0.28,0.05), p=0.154).

This suggests that subicular cells, more so than CA1, are influenced by specific biases in animals’ trajectories, which propagate into the successor features and differentiate them from uniform random walks (Fig.6d; see Supplementary Information Figure S1 for consistent results using symmetric, uniform Gaussian basis features).

### Timescales moving from hippocampal to subicular predictions

The anatomical organisation of the hippocampal formation—with information flowing through CA3 and CA1 to subiculum^17^—suggests a hierarchical processing stream. If each element of this hierarchy successively implements predictions appropriate to its inputs, we hypothesised that subiculum should integrate information over longer temporal horizons than CA1, enabling it to capture extended behavioural patterns and environmental affordances. Within the successor representation framework, such differences in temporal integration would manifest as distinct optimal discount factors (γ) for each region. To test this hypothesis, we systematically varied γ and evaluated which values produced the best RSA fits to subicular versus CA1 recordings in a cell-wise manner.

We found that subicular and CA1 representations were optimally captured by markedly different γ values. Subicular cells were best fit with γ values between 0.99375-0.995 (Fig.7a left), corresponding to effective temporal horizons of 16-20 seconds. In contrast, CA1 representations were optimally modelled with γ values of 0.9-0.95 (Fig.7a right), reflecting substantially shorter temporal horizons of 1-2 seconds (Mann-Whitney U test between SUB and CA1 optimal γ values, U=273019, p<0.001).

**Fig.7.**
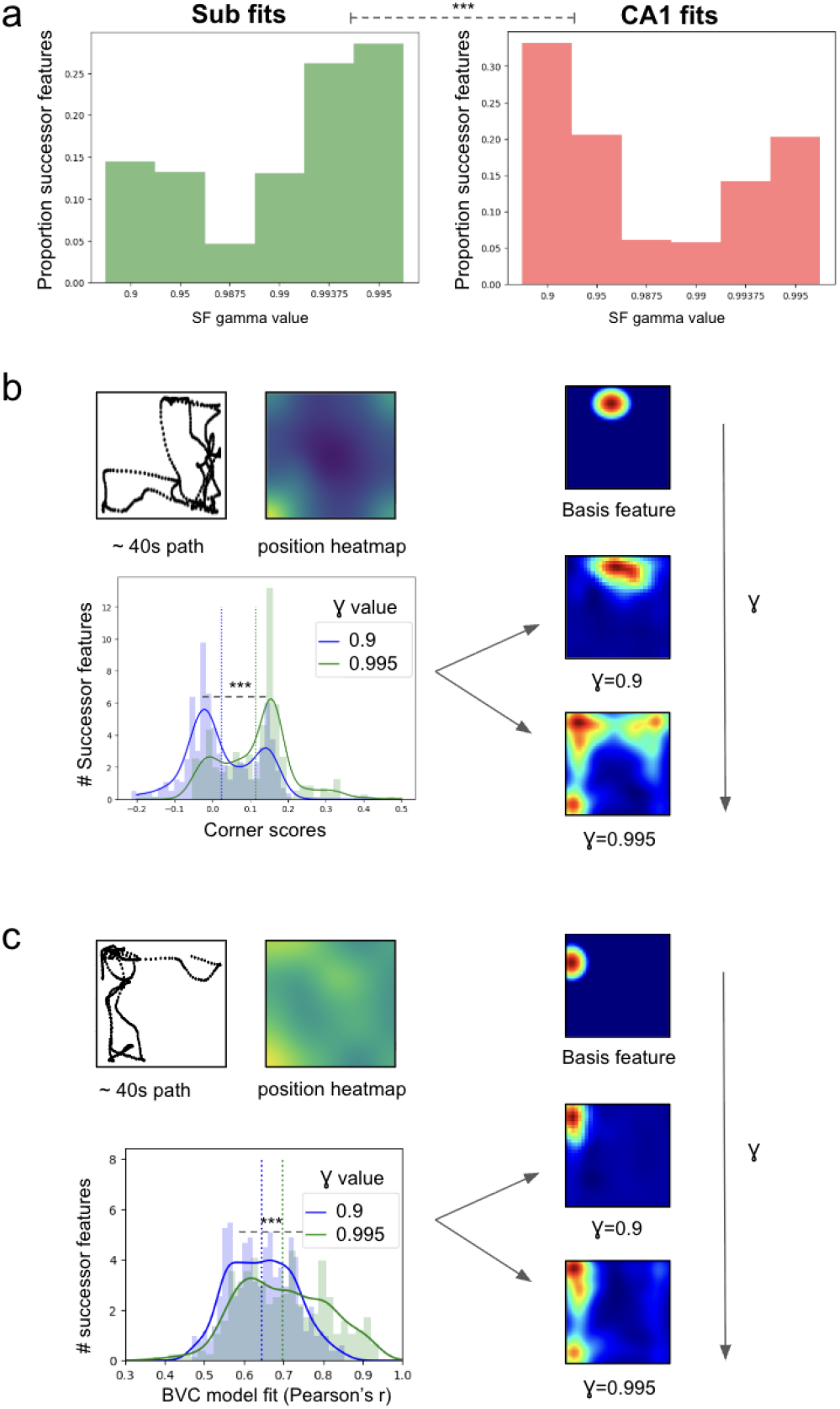
Distinct temporal horizons capture subicular and hippocampal representations. (**a**) Distribution of optimal γ values for fitting subicular (left) versus CA1 (right) cell responses. Subicular cells were best captured with γ≈0.99375-0.995 (temporal horizons ∼16-20s), whilst CA1 representations required γ≈0.9-0.95 (temporal horizons ∼1-2s). (**b**) Transformation of a corner-proximal basis feature with increasing γ. Top: trajectory (∼40s shown) and position heatmap illustrating corner-dwelling behaviour. Right: successor features at γ=0.9 (CA1-like) versus γ=0.995 (subicular-like) demonstrate how extended temporal integration bridges multiple corner locations. Bottom: distribution of corner scores across successor features shifts towards larger values with increased temporal horizons (γ) - vertical lines indicate mean values. (**c**) Emergence of boundary representations with increased temporal horizon. Top: trajectory (∼40s shown) and heatmap demonstrating thigmotaxic behaviour. Right: boundary-proximal basis feature transforms into elongated boundary response at γ=0.995 (∼20s predictive time horizon). Bottom: distribution of boundary vector cell model fits (Pearson’s r) improves with increasing γ, vertical lines indicate mean values. *p<0.05; **p<0.01;***p<0.001.

These distinct predictive horizons have profound consequences for the spatial representations that emerge. To illustrate this, we examined how varying γ transforms a single place cell basis feature. At γ=0.9—characteristic of CA1 processing—the successor feature remained relatively focal, closely resembling its corner-proximal place cell input (Fig.7b right). However, at γ=0.995—typical of subicular processing at 20 seconds—the same corner-proximal basis feature transformed into a response spanning multiple corner locations. Quantitatively, corner scores increased systematically over higher γ values (Fig.7b bottom) (Mann-Whitney U test between corner score values from γ=0.9 to γ=0.995, U=38303, p<0.001).

Similarly, boundary representations emerged preferentially at extended temporal horizons (Fig.7c). When animals executed thigmotaxic runs along walls, successor features with γ=0.995 (∼20s predictive horizon) from boundary-proximal basis features generated elongated boundary fields that captured these extended trajectories. In contrast, γ=0.9 (∼1s predictive horizon) produced more punctate responses that failed to integrate the full extent of wall-running behaviour (Fig.7c right). This transformation was quantified by the progressive improvement in boundary vector cell model fits with increasing γ (Fig.7c bottom), with mean correlations rising from r=0.65 at γ=0.9 to r=0.70 at γ=0.995 (paired t-test, t(399)=7.9, p<0.001)—demonstrating that the diverse spatial phenotypes characteristic of subiculum emerge specifically at extended temporal horizons.

This temporal hierarchy aligns with both the anatomical organisation and functional specialisation of these regions. CA1’s shorter temporal horizon maintains precise spatial coding necessary for navigation and episodic memory formation. Subiculum’s extended temporal horizon enables it to capture behavioural regularities—thigmotaxis, corner-dwelling, trajectory patterns—that unfold over longer periods. Thus, as information flows from CA1 to subiculum, it undergoes a computational transformation from representing immediate spatial states to predicting extended behavioural sequences and environmental affordances.

## Discussion

Here we present a single model of diverse subicular responses, within the framework of predictive representations. We demonstrate that a successor framework trained on biologically plausible CA1 basis features (modelled as place fields) generates spatial responses that closely resemble those reported in rodent subiculum. Specifically, we illustrate that a subset of successor features reproduce boundary vector cell responses^19,20^ and provide a better account of biological boundary-responsive neurons than their namesake model^18^. These cells exhibit behaviour-dependent skewing, as predicted by the successor framework, thus suggesting that the boundary vector cell model might better describe entorhinal border responses^7^ than subicular activity. Our framework also accounts for subicular corner cells^9^, which we show emerge from animals’ idiosyncratic interactions with environmental geometry. At a population level, we demonstrate that the successor framework provides a compelling account of diverse subicular activity across two distinct electrophysiological and one calcium imaging dataset. Our representational similarity analyses reveal that this framework outperforms alternative models in capturing the full spectrum of subicular responses - encompassing both well-characterised cell types and more complex, mixed representations.

Notably, our remapping analyses (Fig.3) reveal that successor representation boundary responses exhibit some resistance to place cell remapping, potentially accounting for the rapid formation and stability of boundary responses observed in novel environments^24^. Similarly, we show that the duplication of boundary responses following barrier insertion^19,20^—a phenomenon considered diagnostic of the boundary vector cell model^18^—emerges naturally when a small subset of place cell fields duplicate against the inserted wall, as documented by Barry *et al*.^19,20^. This finding suggests that the successor framework can account for both the stability and flexibility of subicular boundary representations across environmental manipulations

Thus put simply, the successor representation framework suggests that much of the complexity observed in subicular responses can be reduced to a single predictive objective^38,39^. This can be learnt over states defined by hippocampal place cells via several biologically plausible processes, including spike-timing-dependent plasticity acting over theta sequences^54^, recurrent connectivity^53^, or other forms of synaptic plasticity that do not require theta oscillations^52^, while remaining agnostic to how CA1 place cells themselves emerge.

While our framework accounts for most aspects of subicular responses, certain features warrant further investigation, including the full extent of cross-environment generalisation^8,72^. Although our supplementary results demonstrate that boundary responses can persist despite partial place cell remapping, the complete preservation of boundary vector cell characteristics across entirely different environments^20^ likely involves additional mechanisms. We speculate that subicular networks, with their extensive recurrent connectivity^15,73^, might support compositional successor representation features that can be redeployed across environments. Such a mechanism would complement the stability conferred by non-remapping place cell subsets and could further account for responses that generalise across contexts, such as trajectory dependent firing^23^. Moreover, this combination of stability through preserved place cell inputs and flexibility through recurrent dynamics offers a computational basis for the rapid and flexible adaptation of learned behaviours to novel settings—a key aspect of spatial cognition.

An implication of our work is to propose distinct roles for the two main hippocampal outputs. CA1 appears to transmit a condensed representation of states that are assembled from entorhinal border and grid cells inputs together with path integration and multimodal sensory information, while subiculum represents commonly utilised trajectories afforded by environmental geometry. This differentiation does not negate prior work proposing that CA1 place fields can be understood within a predictive framework^39,45,54,63^. Indeed, Lee and colleagues^63^ demonstrate that when modelling CA1 activity, successor features built on boundary representations (such as those from entorhinal cortex) provide a better account of place cell properties than successor features built on place cell inputs themselves. Our framework extends this, suggesting a multi-level hierarchy where each region computes predictive representations appropriate to its inputs: CA1 builds successor representations over boundary/geometric inputs from entorhinal cortex, whilst subiculum builds successor representations over the resulting place cell representations. This hierarchical predictive map reflects the natural organisation of information flow through the hippocampal-parahippocampal circuit. The key distinction lies in the temporal scope of these predictions—our analyses reveal that subiculum operates with a substantially longer effective time horizon than CA1 (16-20s vs 1-2s). This extended predictive scope enables subiculum to capture complex behavioural patterns and environmental affordances, with CA1 maintains a more immediate representation suitable for precise spatial navigation and memory encoding.

In conclusion, our work provides a unifying computational account of subicular function, positioning it as a predictive map of the environment, derived from hippocampal inputs. This framework not only explains a wide range of observed subicular responses, but also suggests new experimental directions for probing the predictive nature of subicular representations and their role in spatial cognition.

## Methods

### Neural data

We used three experimental datasets in our analysis. Dataset 1 was collected by Muessig and colleagues^21,60,61^ and contains electrophysiological recordings from 11 male (3-6mo) lister hooded rats, freely exploring a plain 62.5×62.5cm square environment for 15 minute trials. Each rat explored the same squared environment for two to three trials per day. The subicular recordings are from three rats^21^ and the CA1 recordings are from a different eight^60,61^. Each rat was implanted with an eight-prong tetrode attached to a micro-drive. All Dataset 1 ratemaps were pre-binned into matrices for each trial. We also obtained the trajectory of each rat for each corresponding trial and environment.

In order to filter out interneurons and axons from CA1 recordings, we applied the triple filter used in Muessig *et al*.^21^, where we selected only ratemaps whose neurons had a mean firing rate less than 5Hz, whose spike width was greater than 0.3ms (peak to trough) and whose mean autocorrelation was less than 25ms. To remove the remaining non-spatial neurons, we only selected cells with a spatial information (SI) score greater than 1. We cleaned the CA1 ratemaps by removing any outer rows or columns with more than five NaNs, a procedure that we repeated twice for each ratemap. We only selected those ratemaps whose resulting shape was 25×25. We then normalised these ratemaps to sum to one and subtracted the minimum value from each entry. We then padded these ratemaps to 27×27 matrices, which resulted in 353 CA1 ratemaps covering two consecutive trials.

Next, we similarly filtered the subicular cells of Dataset 1 to include only those with mean firing rates less than 5 Hz and with a spike width greater than 0.3ms. We removed any outer rows or columns with more than five NaNs and repeated this procedure twice for each ratemap. We then normalised each ratemap to sum to one and subtracted the minimum value from each entry. We padded these ratemaps to 27×27 matrices, which resulted in 335 subicular ratemaps.

Dataset 2 was collected by Sun *et al*.^9^ and includes miniscope calcium imaging data from a total of five male and female mice freely exploring a plain 30×30cm square environment for 20 minute trials. Each mouse explored the same square environment for two trials in the same day, with at least a two hour gap between sessions. We cleaned these ratemaps by removing any outer rows or columns with more than three NaNs, a procedure that we repeated twice for each ratemap. We only selected those ratemaps whose resulting shape was 18×18. We normalised each ratemap to sum to one and subtracted the minimum value from each entry. This resulted in 1937 ratemaps which covered two consecutive trials.

Dataset 3 was collected by Poulter and colleagues^8^ and contains electrophysiological recordings from 6 male Lister hooded rats, aged 3–5 months old at implant, freely exploring a box of size 100×100cm for 20 minute trials. We filtered the pre-binned ratemaps by including only those with a mean firing rate less than 5Hz, as spike width information was not available. We cleaned these ratemaps by removing any outer rows or columns with more than three NaNs, a procedure that we repeated twice for each ratemap. We only selected those ratemaps who had the expected resulting shape of 51×51 matrices. We then normalised each ratemap to sum to one and subtracted the minimum value from each entry. This resulted in 186 ratemaps collected over one trial only.

### Successor features and biologically modelled PC

We generated three separate populations of successor features and biologically modelled basis features using RatinABox^71^ for each of the three neural datasets. First, we created an environment of the same shape as each filtered subiculum dataset (for reference, a 100×100 environment scaled to 1×1 environment). We then selected the longest distance trial trajectory from each dataset’s available rodents and down-sampled it to 10-12 Hz. We used these longest distance trajectories and generated 400 unthresholded, biologically plausible Gaussian basis features for each of the three environments. These basis features followed biologically stable statistics of place cell radii and shapes, with fields being smaller and more elongated near walls, as per Tanni *et al*.^37^:

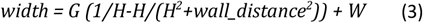

where we set the minimum field width (*W*) to 0.053, wall height (*H*) to 1 and the gain level (*G*) to 0.74 for all successor feature populations. However, we also reproduce key results using uniform and symmetric Gaussian basis features in Supplementary Figure S1. We then normalised each basis feature’s ratemap to sum to 1, subtracted its 40th percentile, and set all negative values to 0.

To calculate the successor features, we thresholded all place cell basis features for ease of training and then used these to update a 100×100 transition matrix, *M*, as per Equation (1). We only updated the matrix, *M*, whenever the agent’s velocity was >0.0001 between timesteps (corresponding to 0.025cm per timestep). We used a learning rate of 0.002 and a gamma of 0.995 for all three populations to optimise convergence. Parameters were initially selected based on standard reinforcement learning values, then validated through systematic parameter sweeps that confirmed distinct optimal regimes for CA1 (γ≈0.9-0.95) versus subicular (γ≈0.995) representations. We generated all successor features using a smoothing sigma of 1.8. We normalised all successor features and thresholded them at the 40th percentile, setting any negative values to 0.

We generated random-walk successor features for each of the three environments using the same 400 basis features, and a trajectory that began randomly within each environment and followed a biologically plausible movement sequence, without thigmotaxis, as defined in RatinABox^71^. We ensured that the average displacement for each random walk was comparable to the average displacement of the rodent for each of the three datasets.

### Boundary vector cell model fits

To fit the boundary vector cell model, we adopted the approach of Muessig *et al*.^21^, utilising an exhaustive search maximising the Pearson r correlation between a candidate rate map and the best fitting boundary vector cell model rate map. Specifically, we generated idealised boundary vector cell model rate maps according to the canonical Hartley *et al*.^18^ model, defining contributions to the boundary response, *g*, as the product of two Gaussians: one tuned to a preferred distance to the boundary, *d*, while the other was tuned to a preferred allocentric direction to the boundary, *ω*.

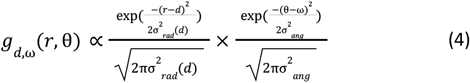

Thus, for a boundary at distance, *r*, and allocentric direction, θ, subtending at an angle, δθ, the firing rate, *f*, of the boundary vector cell is given by:

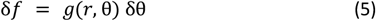

Note, σ_*ang*_ is a constant while σ_*rad*_ varies linearly with preferred distance tuning *d*:

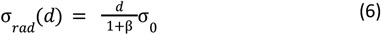

for constant β and σ_0_.

Following Muessig *et al*.^21^, we calculated idealised boundary vector cell firing for each position in a 25×25 unit grid, with constants σ_*ang*_ = 0.2 radians and β=183 units fixed as used in Hartley *et al*.^18^. As with Muessig *et al*.^21^, we generated a total of N=3120 model cells, varying σ_0_ = 6.2, 12.2, 20.2 or 30.2 units, with preferred distance tunings, *d*, ranging from *d*=1 to *d*=13 units in 1 unit increments, and preferred angular tunings, *ω*, varying from *ω=0*° *to ω=354*° in 6° increments.

In order to fit the boundary vector cell model to candidate rate maps (derived from either neural data or our successor representation model), we first resized each rate map, where necessary, in order to fit the 25×25 grid of the modelled boundary vector cells. We then performed resizing by MATLAB R2022b’s ‘imresize’ function with the method set to ‘bilinear’. Next, we performed a Pearson correlation between the candidate rate maps and each of the N=3120 boundary vector cell model ratemaps, identifying the *d, ω*, σ_0_ parameters of the best fitting boundary vector cell. We classified a candidate rate map as a boundary vector cell if the maximum correlation across all model rate map fits exceeded r>0.7. To further analyse the four-fold symmetrical clustering of directional tuning preferences in the square environment, we collapsed direction tunings, *ω*, across all 4 walls, centred on the direction orthogonal to the wall, and quadrupled to span the full 360° before applying circular statistics^74^.

### Remapping analysis

To assess the effect of remapping in place cell basis features (Fig.3a&b), we simulated n=400 randomly spaced basis features from the place cell basis feature model, and trained the successor features using the longest trajectory from the Muessig *et al*.^21^ dataset (see Methods). Upon exposure to the novel environment, a proportion of the basis features (varying from 0 to 1 along increments of 0.1) randomly remapped to new locations. Using these changed basis features, we computed the new successor features relative to the previously learnt successor matrix. By calculating the Pearson correlation between rate maps before and after basis feature remapping, we classified observed remapping in the successor features as r<0.4 and compared this to the same measure calculated on the basis features (Fig.3b), taking the average across 10 random seeds.

To assess the effect of duplication of place cell basis features upon barrier insertion (Fig.3c&d), we simulated n=400 evenly spaced basis features from the place cell basis feature model, and trained the successor features using the longest trajectory from the Muessig *et al*.^21^ dataset (see Methods).

Upon insertion to the novel barrier, a proportion of the basis features with a field on one side of the barrier (varying from 0 to 1 along increments of 0.1) generated a duplicate field on the other side of the barrier. Incorporating these changes to the basis features, we computed the new successor features relative to the previously learnt successor matrix. By calculating the Pearson correlation between the segments of rate maps either side of the barrier post insertion, we classified observed field duplication in the successor features as r>0.4 and compared this to the same measure calculated on the basis features (Fig.3d), taking the average across 10 random seeds.

### Data preparation for behavioural skew analysis

In Figure 4, we analysed the correlation between rodent’s behaviour and the development of asymmetry in subiculum. We performed the following analysis on CA1 neurons from Dataset 1 and subiculum neurons from Datasets 1 and 2, as these were the only two datasets that recorded neurons over consecutive and identical trials. Each ratemap was accompanied by an x/y vector of the rodent’s position over the corresponding trial.

Firstly, we z-scored each ratemap, ignoring NaNs, and set any negative values to 0. We then used scipy’s ‘ndimage’ package^75^ to identify the number of discrete objects in each ratemap. We discarded any ratemaps where we identified more than 5 objects.

To determine the relationship between the behavioural bias of the rodent and the skew of subiculum or CA1 fields in one-dimension, we limited our analysis to those cells whose primary firing field was <=5 to 6 bins of a boundary, where the rodent’s behaviour is more constrained (Fig.2d,e). This corresponds to <12.5.5cm for data from dataset 1 (N=34) and <10cm from dataset 2 (N=113). Dataset 3 did not include consecutive trials without other manipulations. If the number of objects identified was one, we took this object to be the main firing field. If the number was greater than one, we took the field with the highest maximum firing rate of the first three identified as the primary field. We then masked all of the ratemap except for this primary firing field in preparation for the next step.

Next, we chose the axis to collapse the newly masked ratemaps over (horizontal or vertical) as that which gave the lowest maximum when each ratemap was summed over it. Because many fields lay close to a corner and were therefore close to two walls, we only included those cells whose masked ratemaps collapsed over the same axis (horizontal or vertical) in both trials. We calculated the centre of mass of each masked ratemap, now averaged onto one axis, and used the change in this between two trials to quantify the overall change in field skew.

For the behavioural skew analysis, we first isolated the subsections of the animal’s trajectory that ran through each cell’s primary firing field (defined above). We then calculated the resulting allocentric direction between each consecutive position, if that distance was >0.0001 along either the x or y dimension (i.e. the agent moved >0.025cm on either axis). We collected these along the whole trajectory and binned them into a single histogram for each trial, between the allocentric angles of [-π/4, π/4, π3/4, π5/4] as up, right, left and down. We calculated the overall behavioural skew of the animal’s movement through this firing field by subtracting the size of the histogram bar for the ‘backwards’ direction (left or down for fields running along horizontal and vertical walls respectively) from the ‘forwards’ direction (rightwards or upwards for fields running along the horizontal and vertical walls respectively).

Figure 4e,f presents the correlation of the change in the skew of each cell’s main field against the rodent’s change in behavioural bias through that field between the two trials. In order to exclude any cells that are unstable between trials, we only included filtered ratemaps (displaying only the primary field) with at least a 0.3 correlation between trials 1 and 2, however using other thresholds did not significantly alter the results.

### Corner score analysis

For our corner cell analysis, we repeated the procedure of Sun *et al*.^9^ for square, hexagon, triangle and circular-shaped environments. Since it is unclear how Equation 3 can be extrapolated to non-rectangular environments, we generated populations of 100 symmetric Gaussian place cell basis features, with standard deviation 6cm, and calculated the successor features based on the longest distance rodent trajectory for each geometry, as outlined in Section 1. Following Sun and colleagues^9^, we thresholded each successor representation and place cell ratemap at 30-40% of their maximum value to best isolate their main firing fields. Next, we set all values of these ratemaps that were negative to be 0, and performed object labelling using scipy’s ‘ndimage’^75^. For each identified object in each ratemap, we calculated a corner score based on the distances between the object’s centroid, the environment’s centre, *d*1, and the nearest corner, *d*2, as per Sun *et al*.^9^:

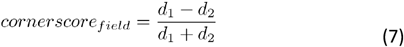

For ratemaps where the number of identified objects was less than the number of corners, we calculated the overall corner score of a ratemap with *k* corners and *n* fields as per Sun *et al*.^9^:

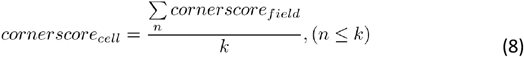

For ratemaps where the number of identified objects was greater than the number of corners, *k*, we calculated the corner score using the top *n* scores as:

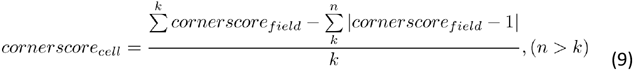

In order to compare the distribution of corner scores between subicular cells and successor features, we generated 2000 shuffled successor features and took the 95th percentile of their corner scores as a benchmark. We calculated these shuffled successor features for each geometry by randomly shuffling the columns of our transition matrix, *M*, after it had been trained using the rodent’s trajectory, and re-generating 2D successor features based on this shuffled matrix. As per Sun *et al*.^9^, we did not include the punishment term for extra firing fields when we calculated the corner scores for shuffled successor features. Figure 5f compares the percentage of successor features and the percentage of place cells that were classified as corner cells by this shuffled threshold, to the percentage of corner cells that are reported for each geometry in Sun *et al*.^9^, using their own spike-train shuffle threshold.

### Relationship between discount factor and time horizon

As shown in George et al.^54^, the definition of successor features can be applied to continuous time by following a temporal discounting scheme determined by an exponential decay with time constant τ:

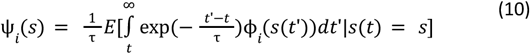

Where *E* denotes the expectation and 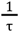 is a scaling constant such that 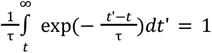.

We can compare this definition to that of the discrete timestep successor features^45,51,76^:

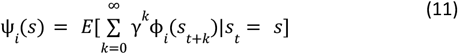

Thus, if consecutive discrete time steps *t* and *t*^′^ increment at a fixed *dt*, such that *t*^′^ = *t* + *dt*, to preserve equivalence between the discrete and continuous definitions we find:

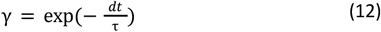

So rearranging for time constant τ yields:

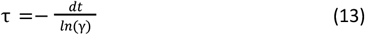

Since the rodent trajectories used in this analyses were downsampled to approximately 10Hz, that sets *dt* ≈ 0. 1 seconds. Thus for γ = 0. 9, τ = 0. 95 seconds, approximately equivalent to a continuous time horizon of 1 second, and for γ = 0. 9, τ = 19. 95 seconds, yielding an equivalent continuous time horizon of approximately 20 seconds.

## Data availability

The neural recordings analysed in this study were obtained from previously published datasets (Muessig *et al*. (2015), Muessig *et al*. (2019), Poulter *et al*. (2021), Muessig *et al*. (2024), Sun *et al*. (2024)). Ownership and responsibility for the dissemination of this data remains with the original authors.

## Code availability

The code required to replicate the analysis in this study is publicly available at: https://github.com/Lauren2909/Unifying-Manuscript.

## Acknowledgements

We would like to thank Dr Daniel Bush and Prof Tim Behrens for providing feedback on the manuscript. This work was supported by the John Monash Foundation (LB) and a Wellcome SRF (CB); BBSRC grants to CB (PI) and CL (co-I) (BB/Y0117751); CL (PI) and TJW (co-I) (BB/M008975/1); and to CL (PI) and SLP (Co-I) (BB/T014768/1). It is also supported by a Wellcome SRF to TJW (220886/Z/20/Z) and an ERC Consolidator to FC (‘DEVMEM’). LMG is an HHMI Investigator and this work was supported by a 1R01MH126904-01A1 (LMG), R01MH130452 (LMG), BRAIN Initiative U19NS118284 (LMG), P50 DA042012 (LMG), The Vallee Foundation (LMG), The James S. McDonnell Foundation (LMG), The Simons Foundation 542987SPI (LMG), NIH K01DA058743 (YS) and The Simons Foundation SCGB Transition to Independence Award (YS).

## Ethics declarations

The authors declare no competing interests.

