## Supplementary Information for "Subicular spatial codes arise from predictive mapping"

##### Additional RSA results

Fig.S1 provides further RSA comparisons with other candidate models. Importantly, subicular cells are also well fit by successor features that are built on symmetrical and uniform Gaussian basis features, which do not deform in size and shape around boundaries. Further, all subicular recordings are significantly less well modelled by both traditional boundary vector cells[1], and uniform, symmetric Gaussian basis features, compared to successor features built on biological basis features and real rodent trajectories.

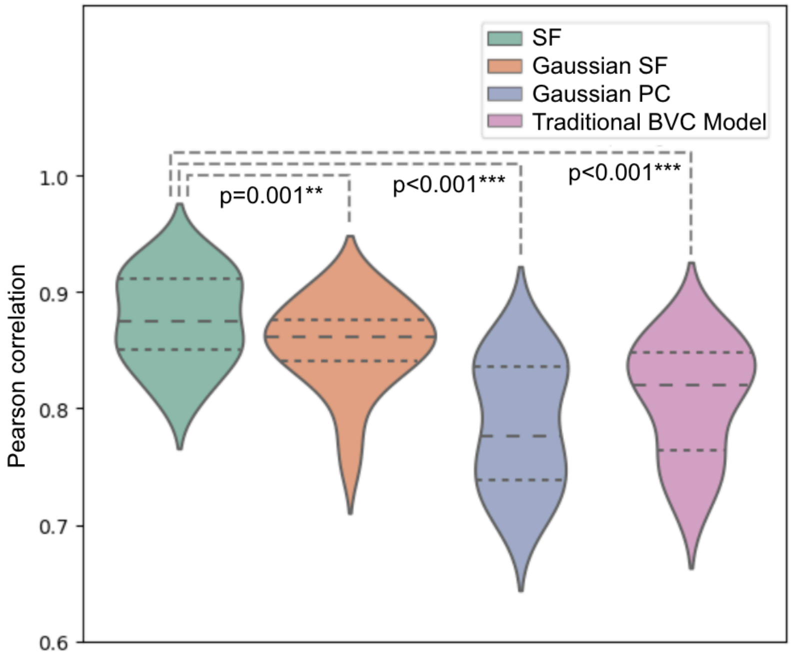


**Fig.S1 Additional RSA correlations to subiculum recordings.** All subicular recordings are better fit by successor features built on biologically-scaled basis features and real rodent trajectories, than successor features built on uniform and symmetric Gaussian place cell bases and real rodent trajectories. However, all subicular recordings are distinctly better fit by both populations of successor features than either Gaussian basis features, or traditional 'boundary vector cells'[1]. *p<0.05; **p<0.01;***p<0.001.

### Appendix

#### Derivation of Successor Feature learning rule

The learning rule (Equations 1-2) is derived using semi-gradient temporal difference learning[2]. Starting with the definition that successor features $\boldsymbol{\psi}(s)$ represent the expected discounted future occupancy of basis features $\boldsymbol{\varphi}(s)$:

$\boldsymbol{\psi}(s) = E[\Sigma\ \gamma^t \boldsymbol{\varphi}(s_t) \mid s_0{=}s]$

Thus,

$\boldsymbol{\psi}(s_t) = \boldsymbol{\varphi}(s_t) + \boldsymbol{\psi}(s_{t+1})$

We seek a matrix M such that $\boldsymbol{\psi}(s) = M\boldsymbol{\varphi}(s)$. Using semi-gradient descent, we minimise square of the temporal difference error: $\boldsymbol{\delta}^2 = [\boldsymbol{\varphi}(s_t) + \gamma\boldsymbol{\psi}(s_{t+1}) - \boldsymbol{\psi}(s_t)]^2$

The key insight is that we only take the gradient with respect to $\psi(s_t)$, treating $\psi(s_{t+1})$ as fixed (hence "semi-gradient"). This yields: $\partial\boldsymbol{\delta}^2/\partial M = -2\boldsymbol{\delta}\boldsymbol{\varphi}(s_t)^T$

Following the negative gradient gives our update rule: $M \leftarrow M + \alpha[\varphi(s_t) + \gamma\psi(s_{t+1}) - \psi(s_t)]\varphi(s_t)^T$
